# Prostacyclin Promotes Degenerative Pathology in a Model of Alzheimer’s disease

**DOI:** 10.1101/2020.04.15.039842

**Authors:** Tasha R. Womack, Craig Vollert, Odochi Nwoko, Monika Schmitt, Sagi Montazari, Tina Beckett, David Mayerich, Michael Paul Murphy, Jason L. Eriksen

## Abstract

Alzheimer’s disease (AD) is a progressive neurodegenerative disorder that is the most common cause of dementia in aged populations. A substantial amount of data demonstrates that chronic neuroinflammation can accelerate neurodegenerative pathologies, while epidemiological and experimental evidence suggests that the use of anti-inflammatory agents may be neuroprotective. In AD, chronic neuroinflammation results in the upregulation of cyclooxygenase and increased production of prostaglandin H_2_, a precursor for many vasoactive prostanoids. While it is well-established that many prostaglandins can modulate the progression of neurodegenerative disorders, the role of prostacyclin (PGI_2_) in the brain is poorly understood. We have conducted studies to assess the effect of elevated prostacyclin biosynthesis in a mouse model of AD. Upregulated prostacyclin expression significantly worsened multiple measures associated with amyloid disease pathologies. Mice overexpressing both amyloid and PGI_2_ exhibited impaired learning and memory and increased anxiety-like behavior compared with non-transgenic and PGI_2_ control mice. PGI_2_ overexpression accelerated the development of amyloid accumulation in the brain and selectively increased the production of soluble amyloid-β 42. PGI_2_ damaged the microvasculature through alterations in vascular length and branching; amyloid expression exacerbated these effects. Our findings demonstrate that chronic prostacyclin expression plays a novel and unexpected role that hastens the development of the AD phenotype.

## Introduction

Alzheimer’s disease is an incurable neurodegenerative disorder that is the most common cause of dementia. The late-onset form of AD is a slowly developing, progressive disorder, with age as the most significant single risk factor. Major pathological hallmarks of the disease include the accumulation of extracellular amyloid protein and intracellular tau protein, accompanied by prominent, widespread neuroinflammation (1–4). With disease progression, numerous neuroinflammatory molecules, including prostaglandins and cytokines, become dramatically upregulated within cerebrospinal fluid and throughout the brain parenchyma, and are associated with cognitive impairment (5–8). These findings suggest that persistent inflammation may be a driver of neurodegenerative disease (9–11). Furthermore, numerous epidemiological studies have suggested that nonsteroidal anti-inflammatory drugs (NSAIDs) may be potentially neuroprotective (12–14); some clinical trials have also reported protective effects of NSAIDs in AD (15–18). In animal models of neurodegenerative disease, numerous experimental studies have demonstrated that NSAIDs have a protective role.

NSAIDs exert their activities through the inhibition of cyclooxygenase (COX) enzymes, which are the mediators of prostanoid biosynthesis. Prostanoids represent a broad class of arachidonic-acid derived molecules, including thromboxane, prostaglandins, and prostacyclin, that can exhibit paracrine signaling activity. Typically, these prostanoids fill a variety of diverse physiological processes that are important for tissue homeostasis (19–21). Inflammatory events and neurotoxic insults can significantly upregulate prostanoid expression, a process that significantly contributes to the AD-associated neuroinflammatory response (22–25). In Alzheimer’s disease, one of the crucial triggers associated with prostanoid up-regulation is the accumulation of amyloid-beta protein, which both stimulates pro-inflammatory cytokine production and increases cellular phospholipase activity, the first step in prostanoid biosynthesis (26–28).

The liberation of fatty acid substrates, such as arachidonic acid, is the initiating step of the prostanoid signaling cascade. Free arachidonic acid undergoes conversion, in a two-step reaction, to PGH2 by COX enzymes. Two major COX enzymes exist, COX-1 and COX-2, which have distinct but overlapping tissue profiles and activities. COX-1 is expressed in the periphery and is a consistently active enzyme that is important for the maintenance of tissue homeostasis. In contrast, COX-2 represents an inducible form of the enzyme. In the brain, COX-2 expression usually is low but becomes dramatically upregulated in Alzheimer’s disease (29, 30). The reported neuroprotective properties of NSAIDs are associated COX-2 (31, 32), and numerous studies support the role of COX-2 upregulation in the pathogenesis of AD. Numerous prostaglandins such as prostaglandin E2 (PGE_2_), prostaglandin D2 (PGD_2_), prostacyclin (PGI_2_), prostaglandin F2α (PGF_2α_), and thromboxane A2 (TXA_2_) become upregulated with COX-2 expression; several of these prostanoids, such as PGE_2_ and PGD_2_, can stimulate amyloid-beta production and can play roles in other amyloid associated pathologies (33–36). However, the impact of increased PGI_2_ in AD is not well understood.

Prostacyclin is synthesized from arachidonic acid through a two-step reaction of the COX and PGI_2_ synthase (PGIS) enzymes. PGI_2_ acts via the G-protein coupled IP receptor to activate adenylyl cyclase and PKA, thereby increasing intracellular cAMP to produce vasodilatory and anti-inflammatory effects (37). While prostacyclin signaling is largely associated with peripheral vasoregulatory activity, multiple cell types in the brain express both PGIS and the IP receptor, including neurons, glia, endothelial cells, and smooth muscle cells (38, 39, 40). These findings suggesting that prostacyclin may act as a modulator of CNS activity.

Experimental applications of stable analogs of PGI_2_ within the brain have shown improvements in vascular functions and recovery from neuronal damage. For example, the application of iloprost, a stable analog of PGI_2_, was able to significantly reduce infarct size after 6 hours of middle cerebral artery occlusion in rabbits (41). While the infusion of TEI-7165, another analog of PGI_2_, was able to rescue hippocampal neurons and improve response latencies in a step-down passive avoidance test in gerbils subjected to forebrain ischemia (42). Additionally, PGI_2_ appears to impact influence behavior and cognitive processing positively; CP-Tg mice contain a modified cyclooxygenase-1 enzyme linked to Prostaglandin E synthase, which increases PGI_2_ production (43–45). In our previous work, we found that increased levels of PGI_2_ improved short-term memory as the CP-Tg mice learned significantly faster in training compared to controls in a contextual fear conditioning test (46).

Several studies have suggested that PGI_2_ may have some impact on neurodegenerative disorders such as Alzheimer’s disease (47, 48). He et al. (49) was able to show that agonist-induced activation of the IP receptor-stimulated production of soluble APPα, a neuroprotective isoform of the amyloid precursor protein, in isolated human microvascular endothelial cells. Wang et al. (50) demonstrated that injections of PGI_2_ into an amyloid precursor protein/presenilin 1 transgenic mouse model increased Aβ levels and proposed this was due to upregulation of the amyloid cleaving proteins, γ-secretase APHlα, and APHlβ, via the PKA/CREB and JNK/c-Jun pathways. γ-secretases are responsible for cleavage of APPβ to produce the cytotoxic Aβ_1-42_ peptides, suggesting that PGI_2_ may regulate amyloid-associated pathology. These data suggest that increased PGI_2_ is likely to play a modulatory role in cognitive function associated with amyloid metabolism.

For this work, we evaluated the effect of PGI_2_ overexpression in a model of neurodegenerative disease. CP-Tg mice were crossed to APdE9 mice, a model of Alzheimer’s disease that develops prominent Aβ pathology and develop spatial memory impairments by 12 months of age (51). APdE9/CP-Tg mice, along with age-matched controls, were subjected to behavioral tests to assess possible changes in cognitive and anxiety-like behaviors. To investigate the impact of prostacyclin overexpression on the amyloid phenotype, we performed Aβ ELISAs on whole brain homogenates. We also investigated prostacyclin-mediated changes to the neurovasculature using immunohistochemical imaging.

## Results

### Impacts on non-cognitive behavior

Analysis of open field data, a test for locomotor activity and anxiety-like behavior, revealed that ambulation, resting, and margin times differed among the mouse lines. CP-Tg mice exhibited significant anxiety-like behavior measured by greater times spent resting and in the margin with less time spent moving (Fig. 1, *p* < 0.05). APdE9 lines showed significant increases in ambulation and reductions in resting times, indicating an anxiolytic-like effect when compared to the CP-Tg or NTg controls (Fig. 1, *p* < 0.05). Prostacyclin overexpression in the APdE9/CP-Tg mice did not affect amyloid-mediated anxiety-like behavior as no differences were observed between the APdE9 mice and APdE9/CP-Tg mice (Fig. 1).

**Figure 1.**
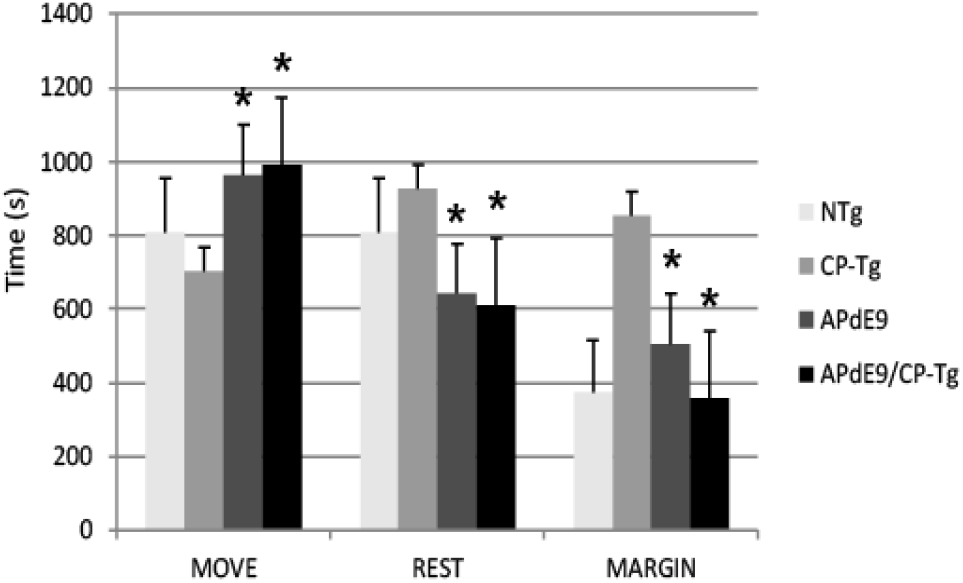
Effects of prostacyclin overexpression in the open-field test in APdE9 and control mice. Increases in ambulation and reduced resting times indicate a reduction in anxiety-like behavior (anxiolytic effect), with the opposite results indicating an increase in anxiety-like behavior (anxiogenic effect). Bars are means ± SEM, n = 7-8 mice per group. *p < 0.05 compared to CP-Tg control mice.

The light-dark exploration test and the elevated plus-maze were used to evaluate anxiety. In the light-dark exploration test, a decrease in exploratory behavior in the lighted area and a preference for the dark compartment was considered a measure of anxiety-like behavior. We found that prostacyclin overexpressing mice spent significantly less time in the light compartment than the NTg control mice (Fig. 2A, *p* < 0.05). Anxiety-like behavior was also measured using an elevated plus-maze, with increased time in the lit open-arm of the plus-maze associated with anxiolytic behavior. Both the amyloid and prostacyclin expressing mice spent significantly less time in the open arms and made fewer transitions, indicative of anxiogenic behavior, compared to the non-transgenic control group (Fig. 2B and 2C, *p* < 0.05).

**Figure 2.**
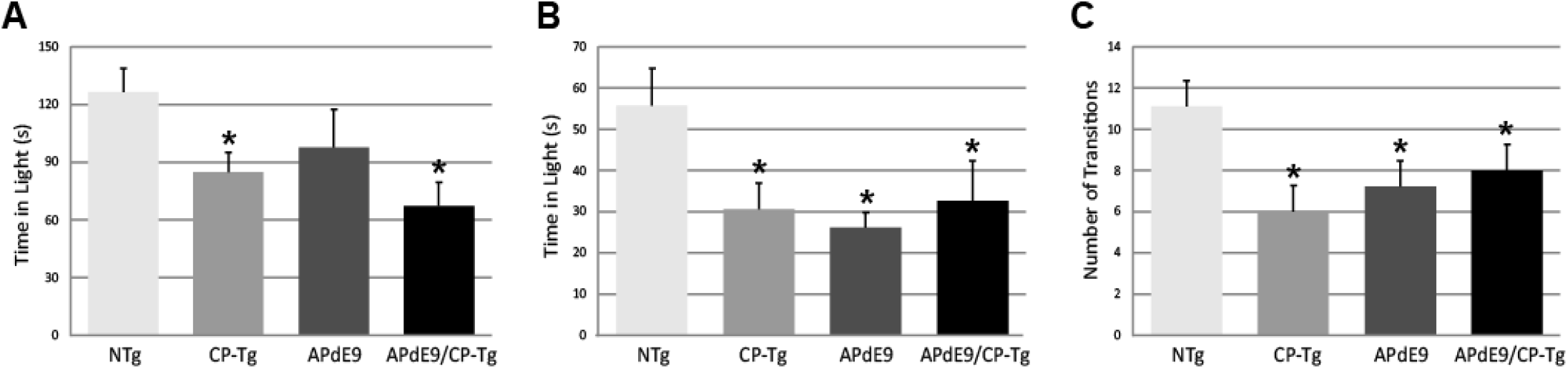
Prostacyclin overexpression increased anxiety-like behavior as measured by light dark and elevated plus-maze tests. (A) CP-Tg and APdE9/CP-Tg mice spent significantly less time in the light during light-dark exploration. (B) In the elevated plus-maze, all transgenic mouse lines spent significantly less time in the lit arms, and (c) and made fewer transitions between the open and closed arms (C) of the elevated plus maze. Bars are means ± SEM, n = 8-10 mice per group. *p < 0.05 compared to NTg mice.

### PGIS overexpression improves motor coordination

Motor function and coordination was assessed using a motorized rotarod. CP-Tg and APdE9/CP-Tg mice had an increased latency to fall in trials 7 and 8 indicating PGIS overexpression increases coordination and balance in mice 6 months of age (Fig. 3, *p* < 0.05).

**Figure 3.**
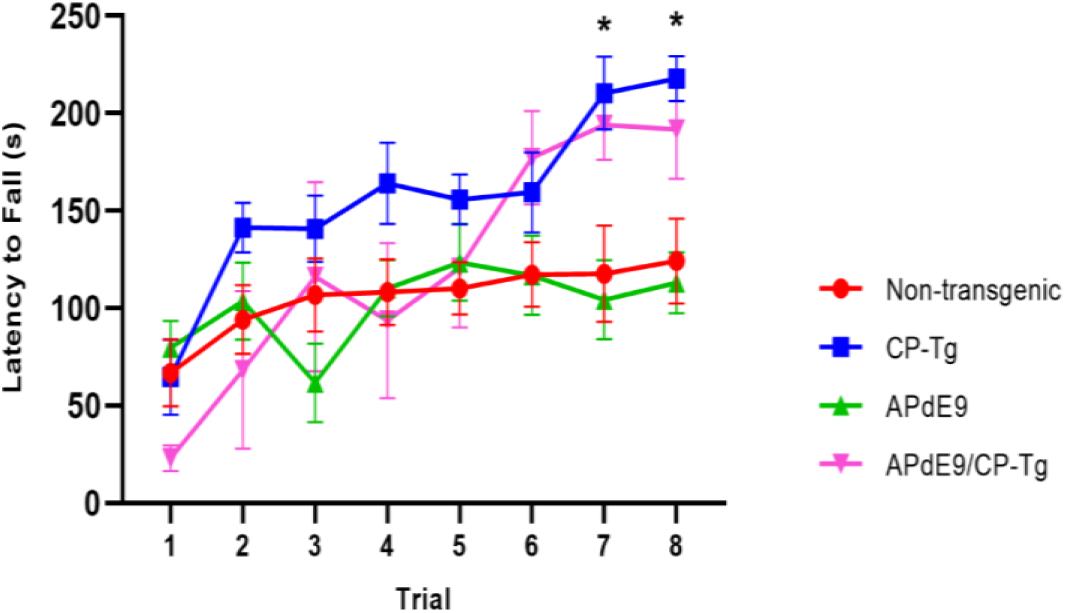
Prostacyclin overexpression enhances motor learning and coordination. In 6month old mice, both CP-Tg and APdE9/CP-Tg mice showed improved coordination on the rotarod, compared with NTg and APdE9 mice on trials 7 and 8. Bars are means ± SEM, n = 7-8 mice per group. *p < 0.05 compared to NTg and APdE9 mice.

### PGIS overexpression impairs associative learning

Context/cue fear conditioning tests were run to assess prostacyclin-mediated changes in associative learning of AD mice measured by percent freezing. During training, mice were conditioned with three shocks paired with a tone. The APdE9/CP-Tg mice displayed delayed learning to the aversive shocks when compared to the APdE9 mice as well as the CP-Tg and NTg controls as the magnitude of the freezing response was decreased in the training period (Fig. 4A, *p* < 0.05). CP-Tg mice also demonstrated delayed learning compared to the APdE9, and NTg controls after the first and second shock but had an increased fear response from that of the APdE9/CP-Tg mice for the 3^rd^ shock in the last two minutes (Fig. 4A, *p* < 0.05). APdE9 mice maintained comparable learning to that of the NTg control mice with no differences seen in percentage freezing during the entire training trial (Fig. 4A).

**Figure 4.**
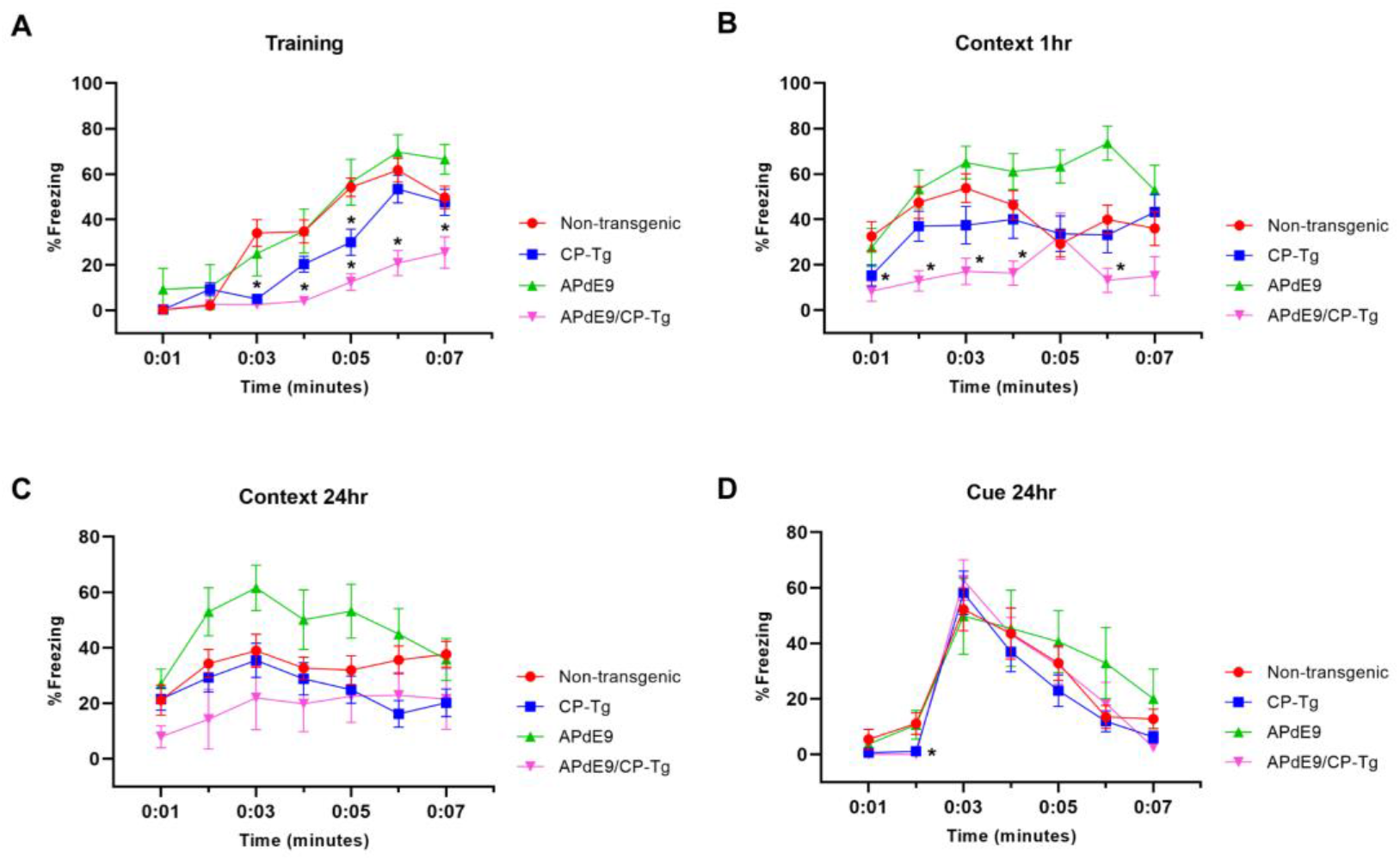
APdE9/CP-Tg mice exhibit impaired learning. (A) PGIS overexpression significantly decreased the percentage freezing time after the first and second shock (shock given at 2^nd^ and 4^th^ minute) of the training trial (p<0.05) while the combination of amyloid and prostacyclin further decreased freezing times during training. (B) APdE9 mice exhibited an enhanced short-term memory response while APdE9/CP-Tg mice exhibited a depressed short-term memory response. (C) CP-Tg and APdE9/CP-Tg mice exhibited worse performance in a contextual test of long term memory compared to APdE9 mice. (D) CP-Tg and APdE9/CP-Tg mice showed an initial delay in freezing at the 2^nd^ minute; however, no significant differences were observed after application of the 3 minute tone from minute 3 to 6 in a cued assessment of long-term memory. N = 7-14 mice per group. *p < 0.05 compared to NTg mice.

A contextual fear conditioning trial was performed 1 hour after training. In the same environment but with no tone presented, APdE9/CP-Tg freezing responses were significantly reduced compared to that of NTg (Fig. 4B, p < 0.05). The APdE9 mice showed a significant enhancement in freezing response during minutes 5 and 6 compared to the control (NTg and CP-Tg) mice (Fig. 4B, p < 0.005).

In order to assess long-term memory consolidation, another contextual trial was performed 24 hours after training. The APdE9 mice maintained a high percentage of freezing times similar to the results seen in the 1-hour context trial (Fig. 4C, p < 0.05). Percentage freezing times for the APdE9 mice were not significantly different from the NTg and CP-Tg controls. During a 24-hour cued conditioning trial where a 3-minute tone is presented in a different environment, all mice exhibited an increased freezing response to the tone with no significant differences by the end of the 7 minute trial period. However, CP-Tg and APdE9/CP-Tg mice did exhibit significantly more acitivity measured by lower percentage freezing, compared with NTg and APdE9 mice, before the tone was presented (Fig. 4D, p < 0.05).

### Prostacyclin drives Aß production and increases amyloid burden

An enzyme-linked immunosorbent assay of Aß40 and Aß42 was used to determine the impact of elevated PGI_2_ synthesis on amyloid production in the APdE9 mouse model. At 17-21 months of age, the APdE9 and APdE9/CP-Tg mice showed significant increases in both PBS-soluble levels of Aß40 and Aß42 compared to the NTg or CP-Tg controls (Fig. 5A, *p* < 0.05). When comparing the APdE9/CP-Tg mice to the APdE9 controls, the double transgenic line exhibited comparable levels of soluble Aß40 and had a selective increase in soluble Aß42 (Fig. 5A, *p* < 0.05). Levels of insoluble Aß40 and Aß42 in the brain were measured using a RIPA extraction. Again, the APdE9 and APdE9/CP-Tg mice had increased levels of both Abeta40 and −42 compared to the NTg or CP-Tg controls (Fig 5B, *p* < 0.0005), with the double transgenic line having an approximately 1.7-fold and 1.5-fold increases in insoluble Abeta40 and 42, respectively, compared to the APdE9 mice.

**Figure 5.**
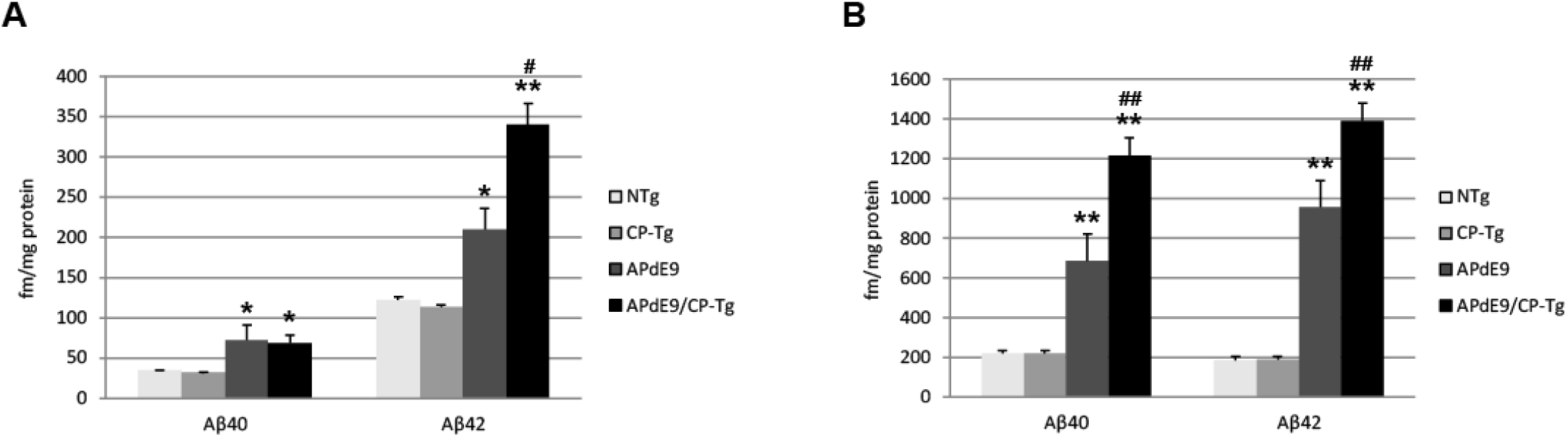
Prostacyclin overexpression increases Aβ production. PBS-(A) and RIPA-soluble (B) Aß40 and Aβ42 levels in APdE9 mice with prostacyclin overexpression. Bars are means ± SEM, n = 10 mice per group. *p < 0.05 and **p < 0.005 compared to control (non-transgenic and CP-Tg) mice. #, p < 0.05 and ##, p < 0.05 compared to APdE9 mice. A two-way ANOVA with a Tukey’s HSD post-hoc was used to determine significant differences.

Increased production was reflected in higher amyloid burdens. Compared to the APdE9 mice, the APdE9/CP-Tg mice contained a significantly increased burden. The average plaque diameter in the APdE9/CP-Tg mice was significantly larger, with a mean 40% increase in plaque volume compared to APdE9 mice, (Fig. 6, *p* < 0.05). Although existing plaques were significantly larger, an notable finding from this study was that there was no significant difference in the number of plaques between different lines.

**Figure 6.**
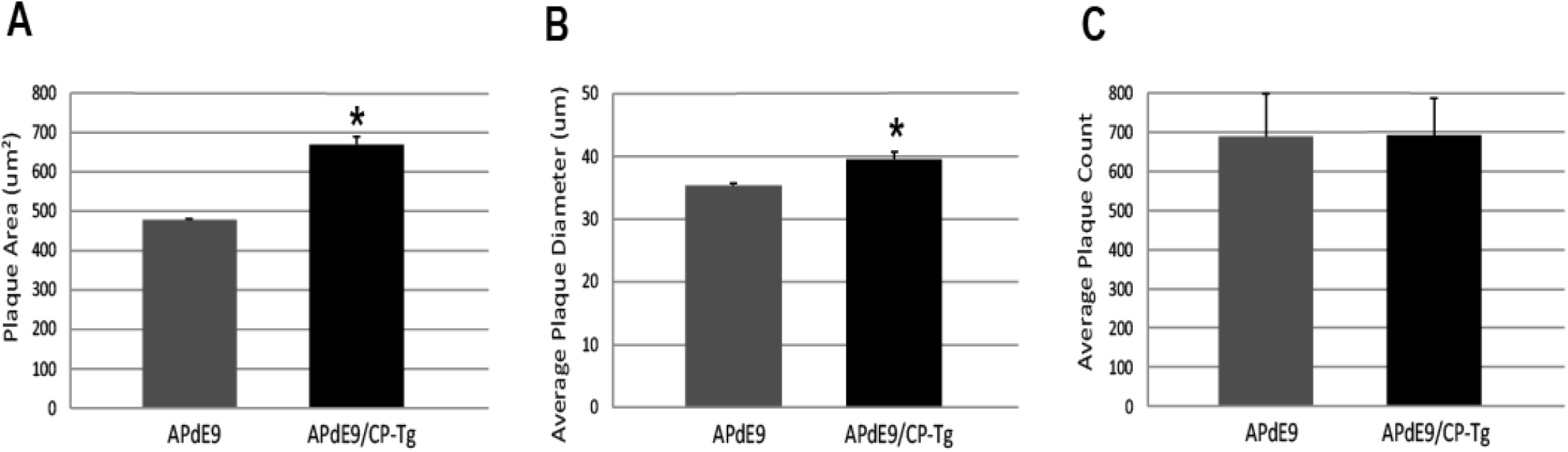
Prostacyclin overexpression increases plaque area and diameter in the brains of APdE9 mice. Immunohistochemistry to detect amyloid plaques (4G8) in coronal brain slices from APdE9 mice (A) and APdE9/CP-Tg mice (B). Scale bar = 200 μm. Measurements in five section of whole cortex determined (C) average plaque area, (D) average plaque diameter (D), and (E) cumulative plaque count per subject. Bars are means ± SEM, n = 10 mice per group. ***p < 0.0005 and *p < 0.05 compared to APdE9 control mice. A one-way ANOVA was used to determine significant differences.

### Loss of pericyte coverage in prostacyclin-overexpressing lines

A colocalization study was performed to assess the effect of prostacyclin overexpression on pericyte death in an AD mouse model. The colocalization of CD-13-positive pericytes with collagen IV-positive microvessels was expressed as Manders Colocalization Coefficients (MCC; M1: the fraction of CD-13 overlapping collagen IV, M2: the fraction of collagen IV overlapping CD-13) (Dunn 2011). In NTg mice, approximately 79% of CD-13 positive pericytes colocated with collagen IV-positive basement membrane, but in CP-Tg mice, this was significantly decreased (*F* (3,95) = 13.41, p < 0.05) to nearly 73% (Table 1). Both amyloid-expressing models exhibited further reductions in pericyte coverage; APdE9 mice had 65% coverage, and APdE9/CP-Tg had 62% (Table 1, p < 0.05). Representative images of each genotype are presented in Figure 7.

**Figure 7.**
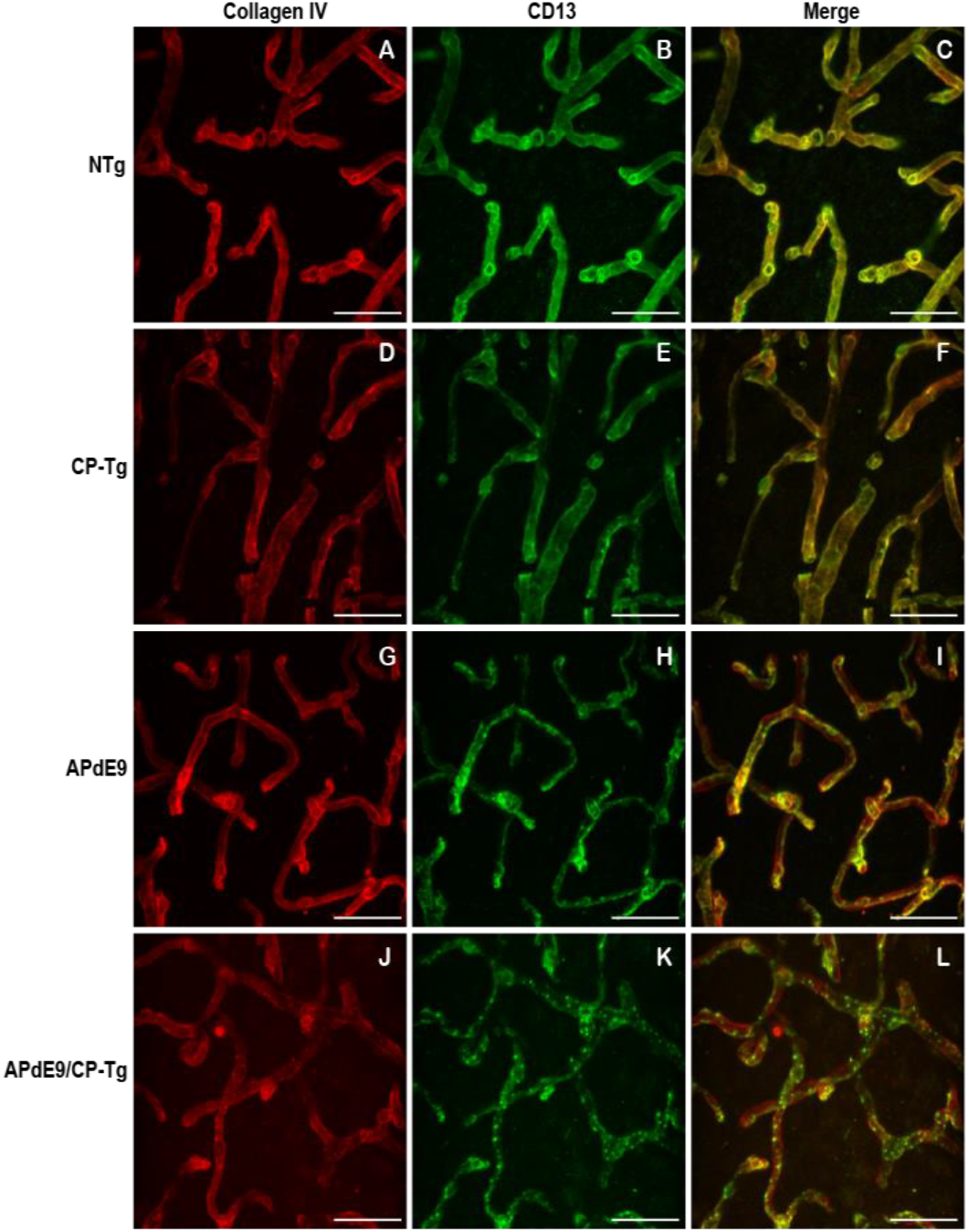
Loss of continuous pericyte coverage in APdE9/CP-Tg mice. 100X confocal image stacks of collagen IV-positive microvessels and CDI3-positive pericytes from the cortex of 17-20-month-old NTg, CP-Tg, APdE9, and APdE9/CP-Tg mice. Scale bar = 30 μm

**Table 1.** Effect of PGIS overexpression on colocalization of microvessels and pericytes. Pearson’s and Manders Colocalization Coefficients (MCC) from dual immunofluorescent stains of collagen IV within the basement membrane of microvessels and CD-13 positive pericytes. M1 signifies the fraction of immunodetectable CD-13-positive pericytes overlapping Collagen IV-positive microvessels. M2 signifies the fraction of immunodetectable collagen IV-positive basement membrane overlapping CD-13-positive pericytes. *p < 0.05 compared to NTg and CP-Tg mice. #p < 0.05 compared to NTg mice.

|  | MCC M1 |  | MCC M2 |  | Pearson's Coefficient |  |
| --- | --- | --- | --- | --- | --- | --- |
|  | Mean | SEM | Mean | SEM | Mean | SEM |
| NTg | 0.808 | 0.012 | 0.788 | 0.020 | 0.859 | 0.013 |
| CP-Tg | 0.796 | 0.016 | #0.728 | 0.022 | 0.774 | 0.019 |
| APdE9 | *0.852 | 0.011 | *0.652 | 0.018 | 0.783 | 0.008 |
| APdE9/CP-Tg | 0.788 | 0.011 | *0.624 | 0.023 | 0.704 | 0.016 |

### APdE9/CP-Tg mice exhibit severe vascular pathology

To examine prostacyclin-mediated structural changes to the cerebral vasculature, we examined stains of the vessel basement membrane using confocal microscopy. 3D image stacks were analyzed using a vessel tracing software plugin in ImageJ. Vascular parameters, including total vessel length, branch number and length, vessel cross-section and diameter, and fractional vessel volume were quantified in the cortex of 17-20-month-old NTg, CP-Tg, APdE9, and APdE9/CP-Tg mice. The vascular parameters are reported as a percent of the total vessel volume imaged. APdE9/CP-Tg mice had significantly shorter and fewer vessels compared to the other models (Fig. 8B-D, p<0.05). CP-Tg mice were found to have a significant increase in both total vessel length and the number of branches compared to NTg mice (Fig, 8B-C, p<0.05). APdE9/CP-Tg mice also presented with the smallest vessel cross-sections and diameters, APdE9 mice with the second smallest and then CP-Tg mice when compared to NTg mice (Fig. 8E-F, p<0.05). The imaged volume fraction occupied by vessels was largest for NTg mice and smallest for APdE9/CP-Tg mice (Fig. 7G, p<0.05).

**Figure 7.**
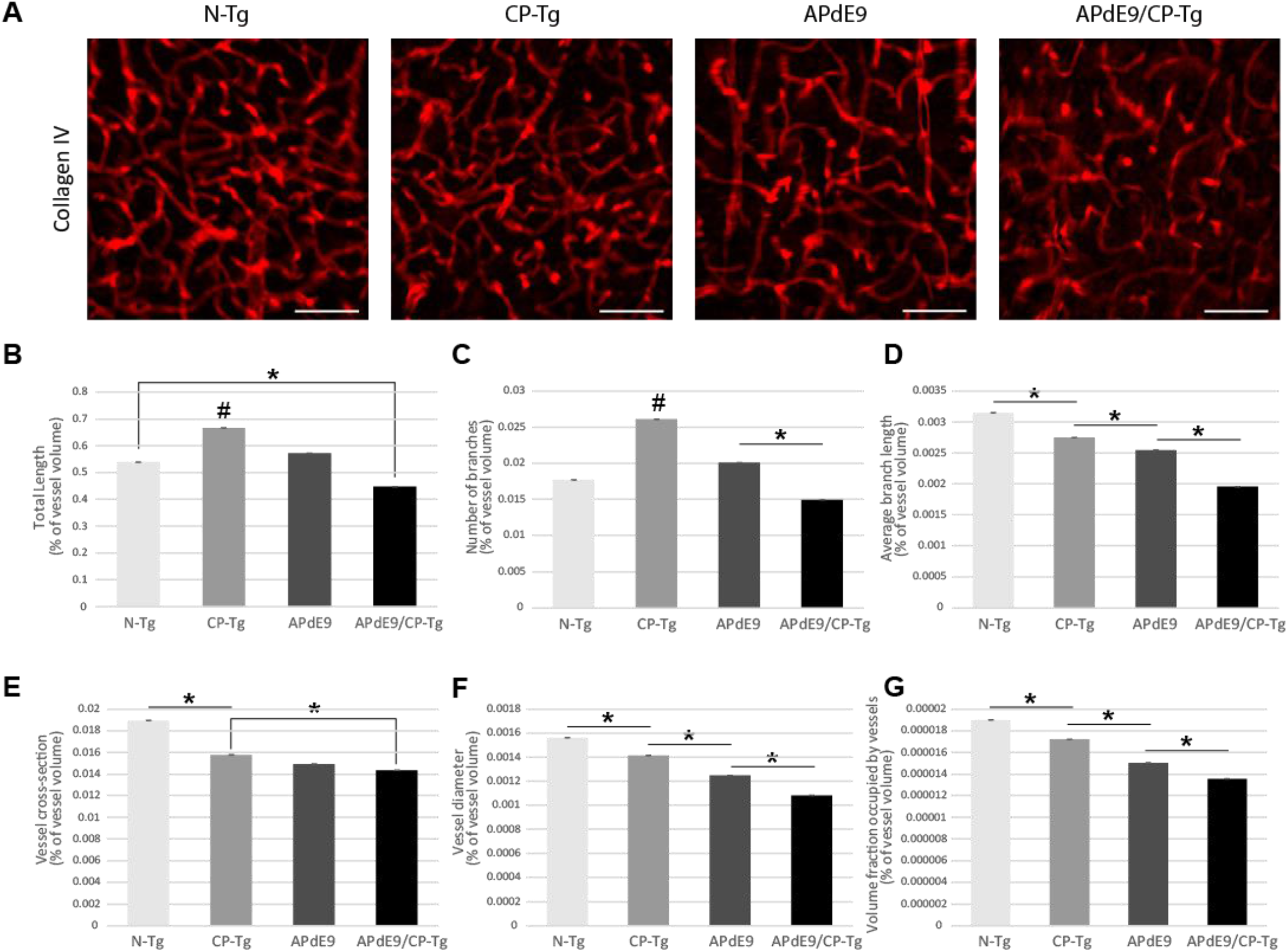
Cerebral vasculature structures are damaged by prostacyclin overexpression in APdE9/CP-Tg mice. Quantification of vascular structures, by collagen IV staining, in the cortex of 17-20-month old NTg, CP-Tg, APdE9, and APdE9/CP-Tg mice. (A) 40X 60 μm thick confocal image stacks of collagen IV-positive microvessels were used for vessel tracing analysis. Scale bar = 50 μm (B) APdE9/CP-Tg mice had significantly fewer vessels compared to NTg or CP-Tg and APdE9 mice, while CP-Tg mice had significantly more. (C) APdE9/CP-Tg mice had significantly fewer number of vessel branches compared to NTg or CP-Tg and APdE9 mice, while CP-Tg mice had significantly more. (D) Branch length is significantly reduced in all models compared to NTg mice; CP-Tg mice show the least reduction in length while APdE9/CP-Tg mice show the greatest. (E-F) Cortical vessel cross-section and diameter were significantly smaller in all models compared to NTg mice with APdE9 mice exhibiting the greatest amount of smaller, constricted vessels. (G) Of the volume of vessels imaged for each genotype, NTg mice maintained the greatest fraction of vessels imaged while APdE9/CP-Tg mice contained the fewest. *p<0.05; #p<0.05 compared to NTg, APdE9, and APdE9/CP-Tg mice. Bars are means ± SEM, n = 5 mice per group.

## Discussion

In this study, we characterized the effect of upregulated PGI_2_ production on behavior, amyloid beta pathology and vessel morphology in a mouse model of AD. Previous work has implicated that exogenous application of PGI_2_ recovers neurological activity in animal models of ischemia or stroke, improves cerebral blood flow, and prevents pericyte loss and vascular leakage after LPS-induced spinal cord injury in mice (41, 42, 59, 60).

We first addressed the possible neuroprotective effects of PGI_2_ overexpression on cognitive and non-cognitive behaviors. In an open field assessment of locomotor activity, both of the amyloid-expressing lines (APdE9 and APdE9/CP-Tg), displayed hyperactivity when compared to the CP-Tg or NTg controls at 12 months of age. CP-Tg mice exhibited decreased activity and spent a larger amount of time along the margin of the field, similar to findings from our previous work (46). Notably, PGI_2_ overexpression, alone and with the APdE9 phenotype, significantly improved coordination by the last two rotarod trials. Elevated plus-maze and light-dark tests were used to assess anxiety-like behavior. Both measures of anxiety-like behavior revealed increased anxiety in all transgenic models (CP-Tg, APdE9, and APdE9/CP-Tg) measured by time spent in the lit compartment and open arms. Previous reports of the APdE9 mouse model observed similar results as the APPswe/PS1dE9 mice displayed increased exploration in an open field and open arms of an elevated plus-maze (51). Notably, PGI_2_ overexpression did not affect anxiety-like behavior in the APdE9 model as the APdE9/CP-Tg mice performed similarly to APdE9 mice. However, PGI_2_ overexpression was sufficient to produce an anxiety phenotype. Recent work has suggested that activation of the IP receptor, a cAMP-dependent PKA pathway, may be sufficient to drive stress responses *in vivo* (61). Stimulation of this pathway leads to activation of CRE (cAMP-responsive elements) binding proteins in the nucleus to synthesize new proteins that alter fear learning and memory formation (61). Increased expression of prostacyclin also appeared to exert detrimental effects on other cognitive tests. We observed impaired learning in APdE9/CP-Tg mice during an associative fear conditioning test. Mice were conditioned to a cue and then assessed for contextual and cued memory to the environment or a tone, respectively. During training, APdE9/CP-Tg mice exhibited a significantly reduced freezing response compared to controls indicating a delay in memory acquisition. Additionally, CP-Tg mice also showed delayed acquisition when compared to non-transgenic controls at the third and fifth minutes following the 2-minute and 4-minute tone-shock pairings. A hippocampal-dependent contextual test was performed 1 hour and 24 hours after training. APdE9/CP-Tg mice again showed reduced freezing response compared to all controls at one hour, and 24 hours were comparable with CP-Tg and non-transgenic controls.

Conversely, previous studies using an exogenous application of a prostacyclin analog, MRE-269, after stroke in aged rats, was able to recover long-term locomotor and somatosensory functions (59). Although IP receptor deletion in mice can be neuroprotective with acute insults such as ischemic damage (62), PGI_2_ overexpression may exert long term adverse effects. Our work demonstrates that elevated activation of IP receptor signaling results in reduced hippocampal memory function in the presence of amyloid-beta insults. Fear behavior during a 24-hour auditory cue test was comparable between all groups. As cued conditioning is hippocampal-independent as it requires the use of the amygdala, our results indicate the combination of amyloid and prostacyclin have little to no effect on amygdala-mediated memory retention.

We also find that PGI_2_ overexpression significantly increases amyloid production in APdE9/CP-Tg mice. ELIZAs of whole-brain homogenates show a more than 1.5-fold increase in insoluble Aß40 and Aß42 in the double transgenic line compared to the APdE9 mice Amyloid burden is directly affect by prostacyclin expression. Intriguingly, these increases are associated with a selective increase of Aß42 in the soluble pool, suggesting that prostacyclin may selectively bias γ-secretase processing. Another interesting observation in this study was the finding that prostacyclin caused an average 40% increase in plaque diameter, but without a detectible increase in the overall number, perhaps suggesting that factors involved in Aß metabolism may be modulated through the IP receptor. These findings are supported by recent work on prostacyclin signaling influencing APP processing mechanisms. Agonist activation of the IP receptor was found to increase the expression of the amyloid precursor protein, resulting in the increased production of Aβ (49). Another report suggests that downstream activation of the PKA/CREB pathway after prostacyclin injections induces upregulation of γ-secretase cleaving enzymes and leads to an accumulation of Aβ c-terminal fragments in an APP/PS1 transgenic mouse model (50).

Substantial evidence has shown that disruptions to the neurovasculature are evident in humans and mouse models of Alzheimer’s disease (AD) that directly coincide with an earlier onset and accelerated progression of AD-related pathologies (63–65). These disruptions include altered cerebrovascular functions such as reduced cerebral blood flow velocities and higher resistance indexes (66) that have been correlated with impaired cognition (67, 68). Structural abnormalities include changes such as capillary atrophy, cell degeneration, loss of tight junction proteins within the blood-brain barrier and basement membrane thickening due to cerebral amyloid angiopathy (CAA) in both mouse models and individuals with AD (69–73). However, a complete description of how vascular pathology can affect AD progression, and the role of the inflammatory pathway in vascular and neuronal injuries, is still being investigated.

Using a volumetric analysis, we measured the impact of PGI_2_ overexpression on vascular structure within the brain, quantifying measures of vessels such as length, branching, diameter, and volume. Our results show that the combination of PGI_2_ overexpression with the amyloid phenotype was quite detrimental and worsened multiple measures of vascular structure. Compared with the control group, APdE9/CP-Tg mice had fewer vessels with shorter, smaller diameters. Compared to NTg mice, APdE9 mice had significantly smaller branch lengths, <u>cross-sections</u>, and diameters, as well as a reduction in total volume. Reductions in microvascular density and vessel constriction have been reported in individuals with AD and mouse models of AD. Other studies have suggested that amyloid beta insults can increase microvascular density in mice (74), and there is evidence of angiogenic vessels, accompanied by higher microvessel density, in the hippocampus of <u>post-mortem</u> tissues of individuals diagnosed with AD (75). Moreover, some studies have found no changes in vessel volume in both humans and APP/PS1 mice (76–78). In our studies, we also found an increase in total vessel length and degree of branching in the CP-Tg mice compared to their non-transgenic controls or both amyloid models. However, the CP-Tg mice exhibited significantly smaller diameters and crosssections and had a lower volume than that of NTg mice, indicating that the overproduction of PGI_2_ is detrimental to the neurovasculature.

The regulatory function of the IP receptor in the vasculature of the brain is poorly understood. Consequently, alterations in vasculature led us to examine perivascular cells or pericytes of the BBB. These cells maintain the integrity of the cerebral vasculature by promoting tight junction protein expression, facilitate cell to cell alignment and are integral to preventing leakage of neurotoxic macromolecules that lead to neuronal damage (79). Application of PGI_2_ after LPS-induced vascular damage in a mouse model of spinal cord injury was able to rescue pericyte loss and concurrent leakage of blood components into the CNS (60). Previous work has reported that accumulation of amyloid β in the neurovasculature of AD individuals was correlated with reduced coverage of pericytes on capillaries in AD brains compared to controls (71). Furthermore, studies of pericyte-deficient, APP^*SW/0*^ (Swedish mutation) mice showed that they had accelerated levels of amyloid-beta deposition, increased tau pathology, and increased neuronal degeneration by nine months of age when compared to AD control mice (80). In our studies, we evaluated the percentage of CD-13-positive pericytes that co-located with the vessels’ basement membrane as a measure of coverage. CP-Tg mice exhibited a significant reduction in pericyte coverage compared to non-transgenic controls, demonstrating that PGI_2_ plays a modulatory role in pericyte function. We identified significant reductions in pericyte coverage of APdE9 and APdE9/CP-Tg mice compared to Ntg and CP-Tg mice. However, no difference was observed between APdE9 and APdE9/CP-Tg mice, indicating that amyloid-dependent reductions in pericyte coverage are unaffected by prostacyclin overproduction.

## Conclusions

Early changes in the Alzheimer’s disease include progressive dysfunction in cerebral hemodynamics, changes in vascular structure, and impaired clearance of amyloid-beta proteins, changes that often occur substantially before cognitive deficits become evident. The persistent upregulation of inflammatory and immune responses have been strongly associated with age-dependent cognitive decline (11).

A variety of inflammatory conditions, such as AD, are associated with upregulated prostanoid production produced through COX enzyme metabolism. Epidemiological evidence suggests a preventative benefit for using nonsteroidal anti-inflammatory drugs (NSAIDs) as the coincidence of individuals taking NSAIDs long-term for rheumatoid arthritis reduces the risk of developing AD (81). NSAIDs are inhibitors of the cyclooxygenase (COX) enzyme pathway. In clinical studies for neurodegenerative disease, the chronic administration of NSAIDs have suggested that they may be of some benefit for Alzheimer’s disease, but the severe side effects associated with long-term use of these agents has limited additional clinical testing (81).

This study evaluated prostacyclin overexpression in a model of Alzheimer’s disease, using in a transgenic line that favors the production of PGI_2_ over other metabolites of the COX cascade. In studies of COX metabolism, there is abundant literature suggesting that that increased production of thromboxanes and prostaglandins primarily contribute to inflammatory, degenerative processes, whereas prostacyclin is thought to decreases inflammation and function in both vasoprotective and neuroprotective roles. Based on these studies, we hypothesized that prostacyclin would be strongly neuroprotective and vasoprotective, and would slow the development, in a model of AD.

Surprisingly, we found that PGI_2_ expression worsened multiple measures associated the degenerative pathology. We observed that PGI_2_ overexpression influences anxiety, possibly due to increases in cAMP activity that has been reported to alter gene expression of proteins involved with the fear response. APdE9 and APdE9/CP-Tg mice exhibited the same level of anxiety as the CP-Tg mice in the LD and EPM behavioral tests suggesting PGI_2_ does not affect anxiety in the AD model. In a fear conditioning test, the APdE9/CP-Tg mice exhibited significantly worse associative memory acquisition and consolidation than APdE9 controls. Although earlier reports using PGI_2_ analogs suggested activation of the IP receptor could be neuroprotective, CP-Tg mice presented with memory deficits when compared to non-transgenic controls. In our studies, increased production of PGI_2_ accelerated amyloidogenesis by more than 50%, significantly increasing the production of soluble and insoluble Aβ peptides. Overexpression of PGI_2_ severely impacted multiple measures of vascular health, a process that was exacerbated with amyloid-beta pathology. Given that prostacyclin is largely protective when expressed in peripheral tissus, our observations suggest that IP signaling is fundamentally different in the cerebrovasculature.

In conclusion, we have provided evidence that chronic production of the PGI_2_ pathway is potentially pathological and may accelerate the development of changes typically associated with the Alzheimer’s disease phenotype. Future studies should explore the mechanism of PGI_2_ signaling through its respective receptor within specific cell types of the CNS to better understand the role PGI_2_ plays in influencing the progression of neurodegenerative disease.

## Materials and Methods

### Animal Model

All experiments were conducted following approved IACUC guidelines, using approved protocols, and mice were housed at the University of Houston Animal Care Facility. Mice were kept in group cages and exposed to a 12-hour light/ 12-hour dark cycle. To develop the APdE9/CP-Tg mouse model, CP-Tg mice, that express a hybrid enzyme complex linking COX-1 to PGIS by an amino acid linker of 10 residues (COX-1-10aa-PGIS) (43–45), were crossed to APdE9 mice, a bigenic model expressing the human APP Swedish mutation and the exon-9-deleted variant of presenilin-1 (dE9) (52, 53). PCR analysis was used to confirm the genotype. Heterozygous mice were used for all studies. Studies composed of a balanced mixture of male and female mice.

### Open Field Activity

The open-field test was used to analyze exploratory behavior within a 60 cm X 40 cm open chamber in normal lighting conditions. Each animal was placed in the center of the apparatus and were given 30 minutes to freely explore the arena. Movement was monitored by a computer-operated system (Optomax, Columbus Instruments; OH) that recorded the time each mouse spent moving, resting or along the margin of the arena.

### Light Dark Exploration

As a measure of anxiety-like behavior a single mouse is placed in an apparatus consisting of a light and dark compartment separated by a single opening and their movements are recorded (46). Mice were subjected to light-dark exploration test to evaluate anxiety-like behavior at three and six months of age. The light-dark box consisted of a light compartment (27 cm × 27 cm × 27 cm) and a dark compartment (27 cm × 18 cm × 27 cm) separated by a partition with a single opening (7 cm × 7 cm) to allow passage between compartments as previously described More time spent in the dark compartment and fewer transitions is considered anxiogenic. Each trial lasted 5 minutes. The number of transitions were measured manually, with the observer blinded to the genotype.

### Elevated Plus-Maze

Anxiety-like behavior can be assessed by utilizing an elevated-plus maze where the mouse has the option to explore two open freely and lit arms, or two arms closed in with blinders. The plus-shaped apparatus is 40 cm above the ground, and mouse movements were recorded by an overhead camera. Time in the light and number of transitions between arms were manually recorded with the observer blinded to the genotype.

### Rotarod

An accelerating cylindrical drum Rotamex Rotarod machine (Columbus Instruments; Columbus, OH) was used to evaluate motor learning and coordination in 12 month old animals. The rotatod consists of horizontal accelerating rods (4-40 rpm) and plastic partitions between each mouse. The mouse is then subjected to 4 trials a day for two days with 15-minute intervals between each trial. A trial ended when the mouse fell off the rod, time elapsed 300 seconds, or the mouse became inverted twice in the same trial without falling.

### Fear Conditioning

Testing for contextualized and cued fear conditioning was followed as previously described in mice at 6 months of age (54, 55). A standard fear conditioning chamber (13 × 10.5 × 13 cm, Med Associates) with 19 metal rods equally spaced on the floor was used to condition the mice. Training consisted of a 7-minute session where a single mouse was subjected to a foot shock (2-sec, 0.75mA) paired with an auditory tone at 120, 240, and 360 seconds. To test for conditioning to contextual cues, the mouse was returned to the chamber 1 hour and 24 hours after the training session. The contextual tests were also 7-minute sessions. However, the shock was not presented. For the 24 hour cue test, mice were returned to the chamber after the surroundings and smell had been altered and during the 7-minute session a 3-minute tone was presented from minute 3 to 6. Freezing behavior was measured using computer software (FreezeFrame, Med Associates/Actimetrics).

### Aβ ELISA

Mice were sacrificed after behavioral testing, and brains were collected. Briefly, hemibrains were homogenized in 1X PBS buffer (pH = 7.4, 1.0 ml/150 mg of tissue) using a PowerMax AHS 200 homogenizer and centrifuged at 14,000xg for 30 min at 4°C. The supernatant was collected and the pellet re-extracted in RIPA buffer (50 Mm Tris-HCl, 150 mM NaCl, 1 % Triton X-100, 0.5% Deoxycholate, 0.1% SDS, 1X PIC, pH = 8.0) and the supernatant collected again. Sandwich ELISAs were performed for monomeric Aβ42 using the antibodies 2.1.3 (Aβ42 end specific, to capture) and Ab9 (human sequence Aβ1-16, to detect). Standards and samples were added to plates after coating the wells with 2.1.3 antibody in PBS and blocking with Synblock (Pierce). Detection antibodies were then applied and developed with TMB reagent (Kirkegaard and Perry Laboratories). The reaction was stopped using 6% *o*-phosphoric acid and read at 450 nm in a BioTek plate reader.

### Quantification of Amyloid Pathology

4G8 antibody (1:500 dilution; Covance, cat no. SIG-39220) was used to detect amyloid pathology in 10 μm thick coronal brain slices. Slices were blocked for endogenous peroxidase using 3% hydrogen peroxide and blocked using 5% serum. Primary antibody was applied and incubated overnight at 4°C followed by a biotinylated secondary goat anti-mouse antibody (Jackson Immunoresearch) incubation for 20 minutes at room temperature and incubation with a Streptavidin/HRP label (Jackson Immunoresearch), followed by visualization with DAB.

Plaque expression in the entire cortex was quantified for each subject. Briefly, five brain sections, equally spaced through the cortex, from each subject were digitally captured and montaged at 10X using on an Olympus DSU system using Neurolucia (Microbrightfield). The entire cortex was outlined, and Image J particle analysis with thresholding was used to quantify total amyloid burden, number and average size for five sections. Studies were performed blinded for the genotype of each subject.

### Immunofluorescence Staining

The primary antibodies used were Collagen IV (1:400 dilution; Cosmo Bio, Catalog Number LSL-LB1403) and CD13 (1:100 dilution; R&D Systems, Catalog Number AF2335). Mice were sacrificed by decapitation, and brains were dissected. Brains were postfixed by immersion in Accustain (Sigma Aldrich). After fixation, brains were cryoprotected in 30% sucrose and cut into 60 μm thick sections with a cryostat. Free-floating coronal brain slices were subjected to a protein block at room temperature for 30 minutes. Slices were then incubated with primary antibodies at their respective concentrations overnight at 4°C, followed by consecutive incubations of biotinylated anti-rabbit secondary antibody (1:200; Jackson Immunoresearch) and biotinylated anti-goat secondary antibody (1:200; Jackson Immunoresearch) both for 1 hour at room temperature. DyLight fluorophores (Jackson Immunoresearch) were added after each secondary incubation.

### Confocal Microscopy Analysis

Quantification of vessel parameters and pericyte coverage was performed on 60 μm microtome brain slices. Five slices evenly spaced between plates 40 and 65 of the Franklin and Paxinos Mouse Brain Atlas (56) were selected from each of five mice in each genotype for immunofluorescence staining. Blood vessels were visualized by collagen IV and pericytes by CD13 immunostaining. Three randomly selected areas of the cortex from each of the five slices were used for analysis. Z-stacks of 60 μm thickness were captured using a Confocal LS microscope (THOR Labs). Quantification of vessel parameters was performed using a blood vessel and network analysis plugin, Tube Analyst, in ImageJ (57). Pericyte coverage analysis was performed using the ImageJ JACoP plugin as previously described (58). All z-stacks were processed with background subtraction and analyzed using automatic thresholding.

### Statistical Analysis

Results for the contextualized and cued fear conditioning assays were analyzed using a repeated measures ANOVA; we analyzed all other data using factorial ANOVA followed by Fisher LSD post hoc comparisons using Statistica (Tibco Software). A *p* value less than 0.05 was considered significant. All values are represented as S.E.M.

## End Matter

### Author Contributions and Notes

T.R.W., O.M, P.M, D.M, and J.L.E designed research; T.R.W, O.N., C.V, S.M., M.S, and T.B performed research, T.R.W., C.V., O.N, T.B., and J.L.E analyzed data; and T.R.W. and J.L.E wrote the paper.

## Acknowledgments

The authors declare no conflict of interest.

